# Rare morph Lake Malawi mbuna cichlids benefit from reduced aggression from con- and hetero-specifics

**DOI:** 10.1101/2021.04.08.439056

**Authors:** Alexandra M. Tyers, Gavan M. Cooke, George F. Turner

**Affiliations:** School of Biological Sciences, Bangor University, Deniol Road, Bangor. Gwynedd. Wales. UK. LL57 2UW; Max Planck Institute for Biology of Ageing, Joseph-Stelzmann-Straße 9B, 50931, Köln

**Keywords:** Malawi, cichlid, blotch polymorphism, aggression, rare morph advantage

## Abstract

Balancing selection is important for the maintenance of polymorphism as it can prevent either fixation of one morph through directional selection or genetic drift, or speciation by disruptive selection. Polychromatism can be maintained if the fitness of alternative morphs depends on the relative frequency in a population. In aggressive species, negative frequency-dependent antagonism can prevent an increase in the frequency of rare morphs as they would only benefit from increased fitness while they are rare. Heterospecific aggression is common in nature and has the potential to contribute to rare morph advantage. Here we carry out field observations and laboratory aggression experiments with mbuna cichlids from Lake Malawi, to investigate the role of con- and heterospecific aggression in the maintenance of polychromatism and identify benefits to rare mores which are likely to result from reduced aggression. Within species we found that males and females bias aggression towards their own morph, adding to the evidence that inherent own-morph aggression biases can contribute to balancing selection. Over-representation of rare morph territory owners may be influenced by two factors; higher tolerance of different morph individuals as neighbours, and ability of rare morphs to spend more time feeding. Reduced aggression to rare morph individuals by heterospecifics may also contribute to rare morph advantage.

## Introduction

Permanent polymorphism, the presence of multiple genetically determined morphological or behavioural phenotypes within a population, is common in nature and indicates some type of selective balance between morphs. Balancing selection is important for the maintenance of polymorphism as it can prevent either fixation of one morph through directional selection or genetic drift, or speciation by disruptive selection (Huxley 1955; Wellenreuther *et al*. 2014; Kim *et al*. 2019). Polychromatism (colour polymorphism) can be maintained if the fitness of alternative morphs differs in time or space in heterogeneous environments, or if the fitness of a phenotype depends on its relative frequency in a population (Hughes *et al*. 2013; Pérez i de Lanuza *et al*. 2017; Surmacki *et al*. 2013; Svensson 2017; Henze *et al*. 2018).

In many taxa, species-recognition cues have diverged through reproductive or antagonistic character displacement to reduce hybridisation or unnecessary exertion and risk of injury among heterospecifics which are not in direct competition for mates or resources (Seehausen & Schluter 2004; Grether *et al*. 2009). Rare colour morphs can benefit from lack of recognition by receiving less mating-related harassment (Takahashi *et al*. 2010) or less intrasexual aggression from conspecifics (Dijkstra *et al*. 2008; Lehtonen 2014; Pérez i de Lanuza G *et al*. 2017; Scali *et al*. 2020). In aggressive species, negative frequency-dependent antagonism, generated through either evolution of an own-morph bias (Dijkstra *et al*. 2008; Lehtonen 2014; Scali *et al*. 2020) or by a dynamic common morph bias based on experience (Bolnick *et al*. 2016), can prevent an increase in the frequency of rare morphs as they would only benefit from increased fitness (due to reduced aggression) while they are rare (Seehausen & Schluter 2004; Dijstra *et al*. 2007; Bolnick *et al*. 2016).

The existence of conspecific aggression biases does not preclude heterospecific aggression completely. Indeed, resent studies suggest that heterospecific aggression as a result of resource competition and reproductive interference may be more common than previously assumed (Grether *et al*. 2009; Drury *et al*. 2020). Regardless of whether heterospecific aggression is due to convergence in territorial signals among species competing for resources or due to misdirection of aggression because closely related species still share similar signals (Losin *et al*. 2016), in a variety of taxa aggression is often higher among more similar coloured than more differently coloured species (Genner *et al*. 1999; Pauers *et al*. 2008; Anderson & Grether 2010; Losin *et al*. 2016). In taxa where multiple ecologically and phenotypically similar species co-exist in the same habitat there is therefore potential for rare morphs to benefit not only from reduced conspecific aggression, but also from reduced heterospecific aggression. A recent study of Midas cichlids, however, demonstrated increased aggression towards rare heterospecific morphs and suggested that this disadvantage may help to explain their lower frequency in natural populations (Lehtonen *et al*. 2015). The role of heterospecific aggression in relation to polychromatism requires further exploration to improve our understanding of how this may contribute to its evolution and maintenance.

The mbuna cichlids of Lake Malawi (and the closely-related ecologically-similar Mbipi of Lake Victoria) provide an excellent system for the investigation of colour polymorphism. Mbuna inhabit densely packed multi-species communities in the shallow-waters and identify conspecific mates and rivals predominantly by their species-specific colour and pattern (*e*.*g*. Seehausen & van Alphen 1998; Couldridge & Alexander 2002; Jordan 2008; Pauers *et al*. 2008). Several species display a polychromatism characterised by the presence of rare “blotched” morph individuals, which occur at different frequencies in different species and populations (Lande *et al*. 2001; Ribbink *et al*. 1983; Konings 2007). While it is likely that predation has played some role in the evolution of this polychromatism (Seehausen *et al*. 1999; Streelman *et al*. 2003; Maan *et al*. 2008), and mate choice may have been involved in the evolution of (partial) sex-linkage (Seehausen *et al*. 1999; Lande *et al*. 2001; Roberts *et al*. 2009), it is thought that intrasexual competition plays a large role in its maintenance (Dijkstra *et al*. 2008; Dijkstra *et al*. 2009b). Although in most species the frequency of rare morphs remains relatively low in all populations, in some, for example *Maylandia callainos* at Thumbi West Island in Lake Malawi, rare morphs can occur with higher frequency, which allows greater ease of observation and collection. Here we used this population to conduct field observations and laboratory behavioural experiments to test alternative hypotheses regarding aggression biases: Do both morphs preferentially direct aggression towards the common (presumably ancestral) morph, or is there an own-morph bias? An own-morph bias could be sufficient to maintain polymorphism through negative frequency-dependent selection, while a common-morph bias would suggest that an additional frequency-dependent process would be necessary to limit an increase in the number of rare morph individuals. We also test for aggression biases towards the common and rare morph from a closely related heterospecific to assess whether this may contribute to balancing selection. We aim to identify potential benefits to rare morphs, which may occur as a result of receiving less aggression, in the natural environment. Additionally, as differences in selection pressures on each sex, due to differences in the type of competition they experience (competition for mates among males and competition for non-mating resources among females) can result in sex differences in the types of aggressive behaviour used during contests (Arnott & Elwood 2009), we also test for sex-differences in aggressive behaviour and aggression biases.

## Methods

### Study system

*Maylandia callainos* (= *Pseudotropheus callainos* or *Metriaclima callainos*) is a member of the ‘mbuna’ complex of rocky shore cichlid fishes endemic to Lake Malawi. Populations of *M. callainos* are found in shallow water, with peak population density between 3 – 10m (full range 0 – 25m). Their natural range is confined to the northern end of Lake Malawi, where they are often found in sympatry with the more widely distributed ecologically similar congeneric *M. zebra*. However, due to human mediated translocations, they are also found in some southern areas. Phenotypes of common and rare morph mbuna differ between species, but within populations, the common morph is often BB (black vertical melanin bars on a blue/dark background) or solid blue/dark body colour, while rarer morphs have a disrupted melanin pattern of many or few blotches/spots on a light (orange/pink/white) body. Blotch polychromatism is not present in all *M. callainos* and *M. zebra* populations; at some localities only the plain blue (B) and BB morph are found, whereas at others, these common morphs may occur along side rare white (W) and orange-blotch (OB) and very rare white-blotch (WB) and orange (O) morphs. In this study we focus on a well established translocated population of *M. callainos* at Thumbi West Island in the Lake Malawi National Park in the southwest arm of the lake which has both B and W morphs. The likely source population of the *M. callainos* at Thumbi West is Nkhata Bay, where they co-occur with a population of *M. zebra* comprised of BB, OB and O morph individuals (fig. 1).

**Figure 1.**
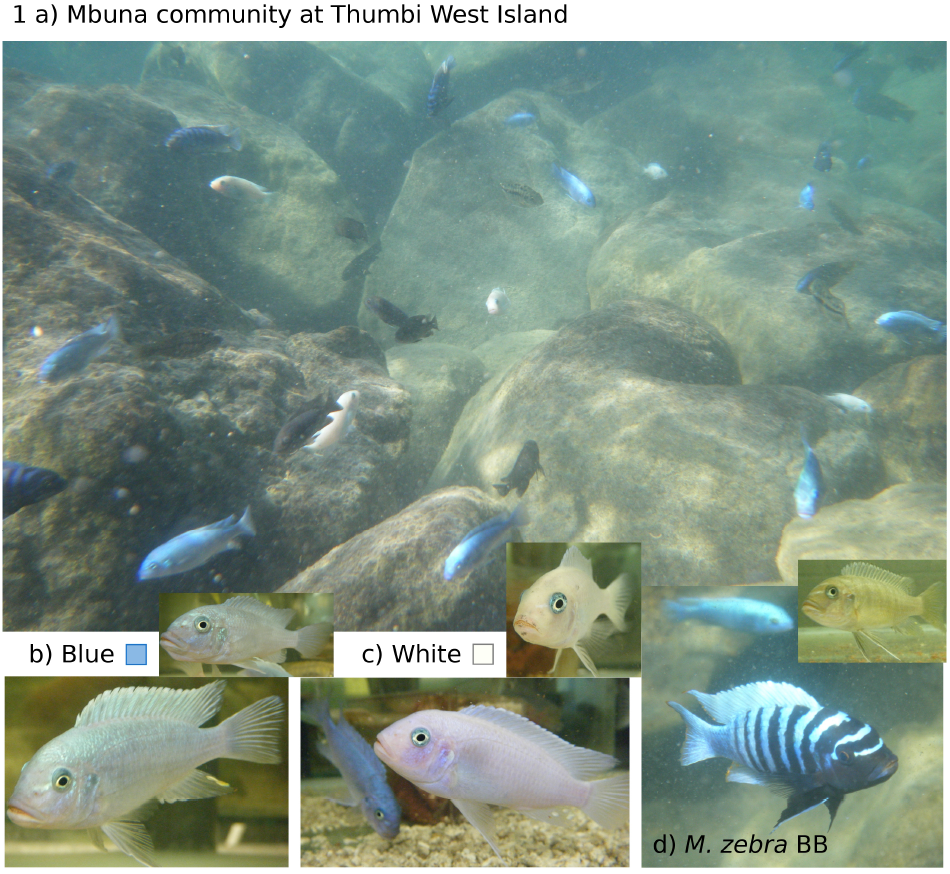
**a)** Mbuna community at Thumbi West Island. Territorial male (bottom) and female (top) of **b)** *M. callainos* Blue morph, **c)** *M. callainos* White morph, **d)** *M. zebra* BB morph. Coloured squares correspond to colours used for the different morphs in results plots.

All fish were wild caught: *M. callainos* and *M. zebra* from Thumbi West Island (TW) in July 2010, *M. zebra* from Nkhata Bay (NB) and Chiofu Bay (CB - naïve to *M. callainos* in the wild and lab) in 2009. Males and females were used in this study, partially because of the lower number of rare males, but also because both male and female aggression biases have previously been suggested to be important in colour polymorphism maintenance in cichlids (*e*.*g*. Dijkstra *et al*. 2008). Furthermore, unlike many species with blotch polychromatism, this one is less strongly female limited, as numerous white *M. callainos* males were found at the study/collection site.

### Field observations (excluding aggression)

#### Frequency of blue and white morph *M*. *callainos* at Thumbi West Island

Snorkel observations were used to estimate the ratio of B to W *M. callainos* morphs in the general population. Dominant mature adult males can be easily recognised by their behaviour and colour, but females and immature males are indistinguishable and are referred to as ‘apparent females’. Hence, the number of males and apparent females of each morph was counted along three 30m transects covering an area half a meter each side of the line (n = 74 fish). The numbers of territory-holding males of each morph were counted in nine 5m2 quadrats (n = 142 fish). Although ideally the comparison should be made between non-territorial males and territorial males, in practice this was not possible due to the difficulty in sexing fish without catching them. However, it is likely that in the whole population, rare morph males occur at a lower frequency than rare morph females (as found in other closely related species with blotch polychromatism, Lande *et al*. 2001; Maan & Sefc 2013), which would make estimates of the ratio of rare to common morph males among non-territorial fish a conservative estimate; territorial W males would be present at a much lower frequency than predicted from the ratio of W morph in the general population.

#### Territory distances between morphs

Territory maps were constructed by drawing the rocky substrate, within 5×5m string quadrats (n = 9), on dive slates while snorkelling. Males frequently return to their spawning cave and focal observations allowed for accurate determination of the position of this territorial focal point for each male within the quadrats. The distance between the territory focal point of each male within the centre 3m^2^ (n = 27 B & 25 W) and closest white and blue neighbour (including fish nearer the edge of the quadrat) was then measured.

#### Grazing differences between morphs

Each grazing action performed by focal individuals was recorded during ten minute observations of territorial males and non-territorial fish (n = 9 individuals of each morph for each social status).

### Field observations of aggressive interactions

During focal observation lasting 10 minutes per fish (n = 9 territorial males of each morph) all aggressive behaviors directed towards the two conspecific morphs were recorded. The vast majority of all aggressive acts recorded were ‘chases’, lateral displays were observed but rare, counts of each type of behaviour were summed for analysis. While collecting data on conspecific aggression biases, aggression towards each focal fish from heterospecifics was also recorded.

### Laboratory aggression trials

To test whether there are differences in the level aggression received by blue and white morph *M. callainos* from conspecifics and heterospecifics, three experiments were carried out using the same methods. Five minute pairwise aggression trials were conducted in two replicate tanks measuring 0.9×0.3×0.3m. Each tank contained a central brick refuge to act as a territory focal point, two transparent (perforated) plastic jars to hold the stimulus fish, an air driven box filter and an internal heater to maintain water temperature at *ca*. 22-24°C. Lights were kept on a 12:12 light:dark cycle. All fish were fed flake food once a day. Females and males were used, but stimulus fish were always the same sex as focal fish. Focal fish were allowed at least 24h to acclimatise before introduction of the stimuli and recording of focal fish behaviour began after emergence from the central refuge. Individual aggressive behaviours (frontal/lateral display, quiver, lunge/butt and bite) were recorded and combined to give an overall aggression count for each individual. To control for potential tank side bias, two separate trials were carried out with each focal fish, each with a different stimulus fish pair and with morphs swapped between sides. To avoid pseudoreplication from the re-use of focal males, before analysis an average was taken of the aggressive behaviour observed in the two trials by each individual.

### Exp. 1: Interspecific aggression biases between species

Firstly, conspecific aggression bias was confirmed by presenting BB *M. zebra* males from CB (n = 10) with pairs of conspecifc BB and heterospecifc B stimulus fish.

### Exp. 2: Intraspecific aggression biases between morphs

For this experiment all available *M. callainos* were used as focal and stimulus fish (n = 10 B male, 6 B female, 3 W male and 9 W female) to test for morph-specific aggression biases among conspecifics.

### Exp. 3: Interspecifc aggression to different morphs

BB *M. zebra* focal fish from different populations (n = 12 male/ 12 female “TW”, 12 male/ 5 female “NB”, 12 male/ 12 female “CB”) were used to test for heterospecific aggression biases to B and W *M. callainos* stimulus pairs. Stimulus pairs consisted of the same *M. callainos* used in Exp. 2.

### Data analysis

Statistical analysis and plotting was carried out using Rstudio (v. 1.2.5033; Rstudio Team 2019) using additional packages Rmisc, pscl, ggplot2, scales. General and generalized linear models were used depending on whether the data originated from continuous measurements (territory distance), or counts (grazing and aggressive behaviour). For the laboratory aggression experiments, trials were omitted from the analysis if the average behaviour count was less than 10. In experiments where females and males were tested, sex was included in the models to test whether this was a significant factor affecting aggression biases towards the stimulus morphs.

#### Frequency of blue and white morphs at Thumbi West Island

A G-test was used to compare the actual number of territory holding W males observed with what would be expected given the proportions of B and W morphs in the general population.

#### Territory distances between morphs

GLMs were used to test: 1) Whether there is a significant difference in the distance between focal fish and the nearest neighbour of the same and different morph, and 2) differences in the average distance to B and W neighbours from focal fish of the two different morphs.

#### Grazing differences between morphs

GLMs were used to test for effects of dominance status and morph on grazing frequency.

#### Field observations of aggressive behavior

Due to the small number of aggressive behaviors and relatively high number of zero counts recorded during observations of aggressive encounters in the field, standard poisson regression GLMs were compared with models corrected for zero-inflation. In most cases the zero inflated model was not significantly better, the results reported here are therefore from the standard Poisson GLMs testing: 1) aggression directed towards territorial intruders of each morph by territorial males of each morph, 2) aggression received by territorial males of each morph from heterospecifics.

#### Laboratory aggression trials

First, GLMs were used first to test whether focal species and sex had a significant influence on the total number of aggressive behaviours (overall aggressiveness of species and sexes) displayed to both stimulus fish. To control for the effect of overall differences in level of aggression between species/populations/sexes/individuals, counts of aggressive behaviour directed towards each of the paired stimulus fish was converted to proportion of aggression. Subsequently, the following were tested: 1) whether *M. zebra* display a conspecific aggression bias when presented with conspecific and common morph heterospecific, 2) whether among conspecifics (*M. callainos*) overall one morph receives more aggression than the other, and whether aggression bias differs among morphs and sexes, 3) whether *M. zebra* display an aggression bias when presented with pairs of common and rare morphs of a heterospecific, and if this aggression bias differs between allopatric populations of *M. zebra*, depending on whether they co-occur with *M. callainos* or not.

## Results

### Field observations (excluding aggression)

#### Frequency of the blue and white morphs at Thumbi West Island

There were significantly (G-test: *G*_1_ = 6.91, p = 0.009) more territory holding rare white (W) morph males than would be expected given the proportion of W and blue (B) morph fish in the general population (fig. 2a).

**Figure 2.**
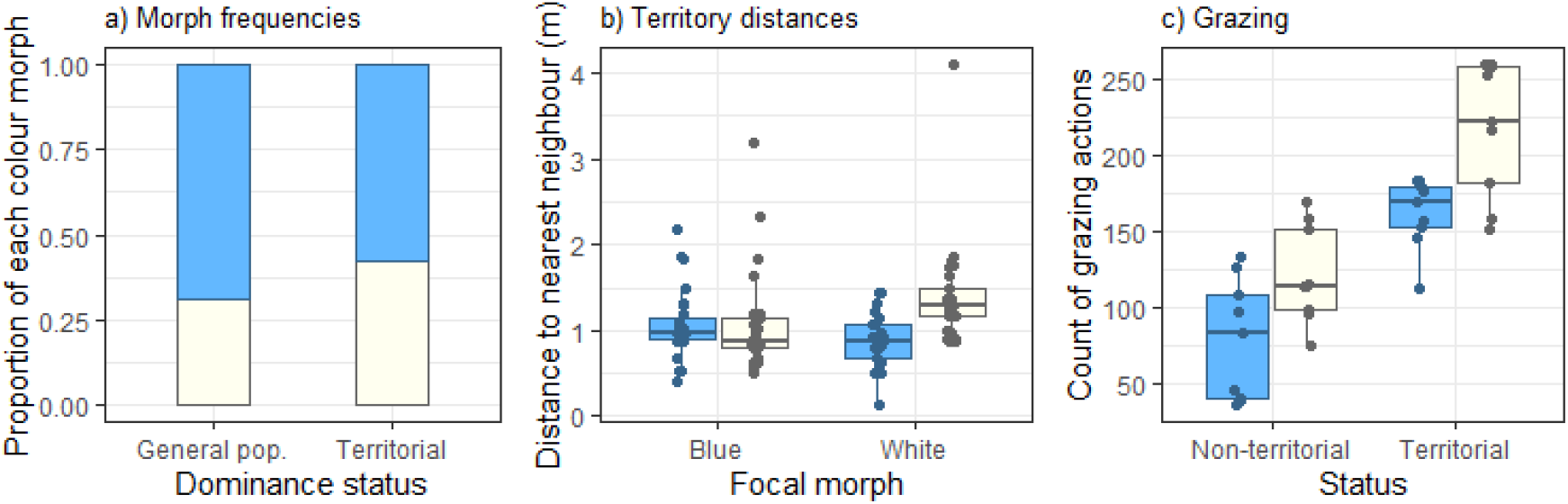
Field observations at Thumbi West Island: **a)** There are significantly (p = 0.009) more territory holding W males than would be expected given the proportion of each morph in the general population. Bars show relative frequency of each morph, n = 216 fish. **b)** Bar colours indicate neighbour colour. Differences in the distance of territory focal point (n = 27 B and 25 W males) among same and different morph males do not always reflect the expectations based on the relative abundance of each in the population. Overall, same morph males hold territories further apart from each other than different morph males (p = 0.0135). The average distance between B males and the nearest B neighbour is no different from between B and nearest W neighbour (p = 0.994), and the distance between W territorial males and their nearest B neighbour is on average the smallest distance recorded between territorial males, despite the lower frequency of W males. **c)** Bar colour indicate focal morph, n = 9 individuals of each morph and status; both territorial and non-territorial W morph fish grazed significantly more than B morph fish (Morph p < 2e-16, Status p < 2e-16).

#### Territory distances between morphs

Firstly, there is a significant difference in the average distance between same and different morph territorial males (glm: t_1,103_ = 2.51, p = 0.0135). On average, territorial males of the same morph are found at greater distance from each other (mean 1.24m) than territorial males of different morphs (mean 0.98m). While it is not surprising to find that W and W are found furthest apart (mean 1.40m), as this would be the assumption based on the observation of a lower frequency of W morph territorial males, if distance between morphs was only based on frequency, it would also be expected that the distance between B morph males should on average be the smallest distance. This is not the case: the distance between B and nearest B is on average the same (mean 1.07m) as, and not significantly different (glm: t_1,53_ = -0.008, p = 0.994) from, the distance between B and nearest W. We also found that the distance between W males and their nearest B neighbour is on average the smallest distance recorded between territorial males (mean 0.89m), and significantly different from the distance between W territorial males and their nearest W neighbour (glm: t_1,49_ = 3.73, p = 0.0005) (fig. 2b).

#### Grazing differences between morphs

Both dominance status and morph had a significant effect on grazing frequency: Compared to territorial males, non-territorial fish grazed significantly more, and regardless of social status B morph fish grazed significantly less than W morph (glm; z_1,35_ Morph = 22.07, p <2e-16, Status = 12.18, p<2e-16) (fig. 2c).

### Field observations of aggressive interactions

Overall, B morph *M. callainos* territorial intruders receive significantly more aggression than their W counterparts (glm; z_1,35_ = -2.79, p = 0.005). However, aggression bias appears to differ between the morphs; B males make significantly more attacks to other B males (glm; z_1,17_ = -2.93, p = 0.003), whereas W morph males show no significant aggression bias (glm; z_1,17_ = 0.69, p = 0.493) (fig. 3a). Although there was a trend towards B fish receiving more aggression from heterospecifics than W fish, this difference was not significant (mean B = 1.8, W = 0.8, z_1,17_ = -1.82, p = 0.068) (fig. 3b).

**Figure 3.**
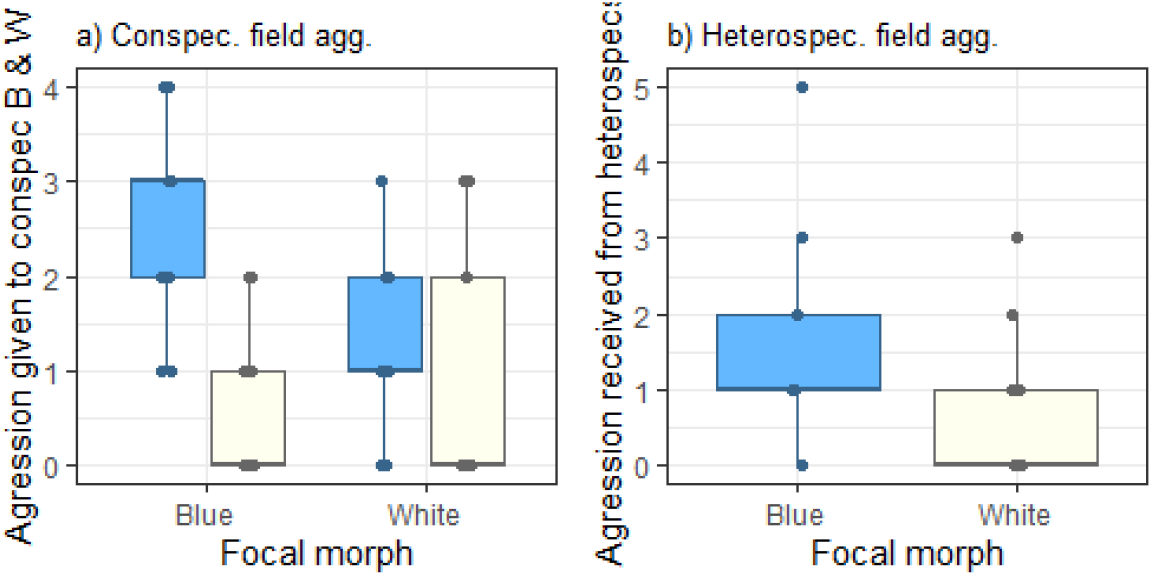
Observation of conspecific and heterospecific aggressive interactions among territorial males in the field: **a)** Bar colour indicates intruder colour, n = 9 focal males of each morph. B morph *M. callainos* territorial males show significantly more aggression towards other B males than towards W males (p = 0.003), W males show no significant aggression bias (p = 0.493). Overall, B morph males receive significantly more aggression (p = 0.005). **b)** Bar colour indicates focal fish colour. There is a non-significant trend towards B morph males also receiving more aggression from heterospecifics (p = 0.068).

### Laboratory aggression trials

#### Differences in focal fish behaviour between species(/experiment) and sex

Within species, both sexes show similar levels of aggression. *M. callainos* overall were more aggressive (glm; Focal.spec z = -10.36, p <2e-16, Sex z = 0.08, p = 0.939, fig. 4a). To test whether this difference in aggressiveness was a real difference between species, or due to *M. callainos* being presented more often with the possibility of being aggressive towards conspecific fish in these experiments, a subset of *M. zebra* CB and *M. callainos* males from the three experiments was compared. *M. zebra* displayed a significantly higher level of aggression in the experiment where the stimulus pair consisted of one conspecific and one heterospecific (Exp. 1) compared to the experiment where they were presented with two heterospecific stimulus fish (Exp. 3). There was no significant difference, however, in the level of aggression between the species in the experiments in which the stimulus pairs contained one conspecific (Exp. 1: *M. zebra* focal fish) or two conspecifics (Exp. 2: *M. callainos* focal fish) stimulus fish (fig. 4b). This suggests that the presence or absence of a conspecific stimulus fish contributed to the overall difference in the level of aggression observed between the species in these experiments. There was therefore no evidence of species differences in intrinsic level of aggression, rather that aggression among heterospecifics is lower than among conspecifics. The attack:display ratio differed between sexes, but not species: Females attack more frequently and males display more (glm; Focal.spec z = 1.72, p = 0.089, Sex z = -4.09, p = 9.45e-05, fig. 4c).

**Figure 4.**
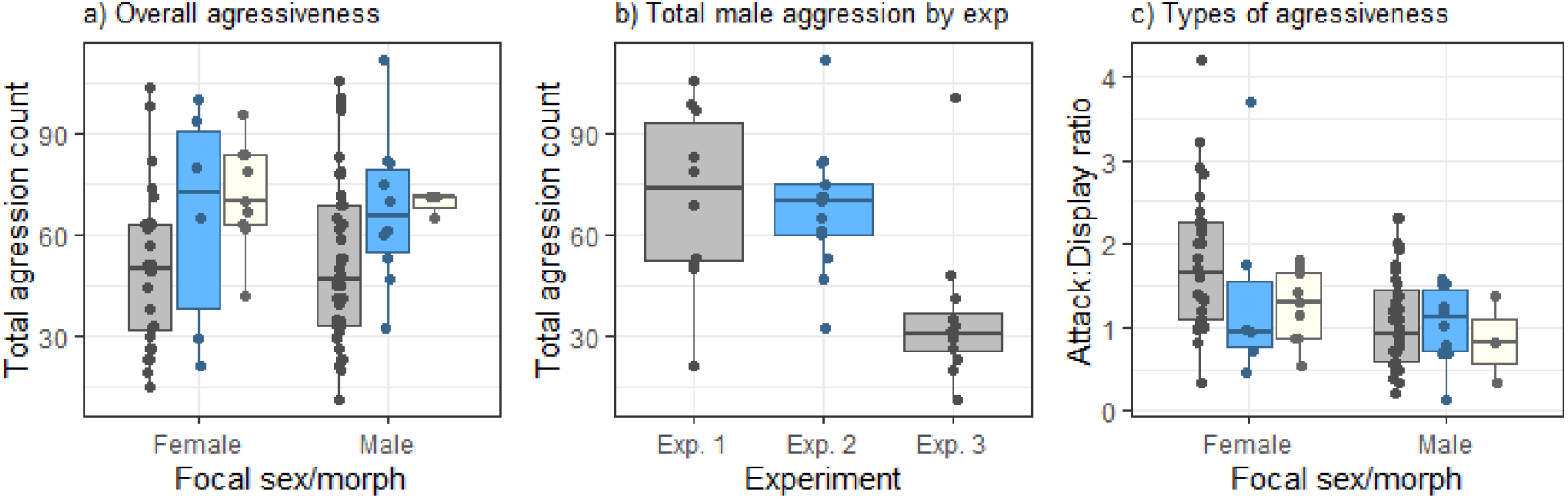
Differences in levels of aggression (total aggression count) and types of aggressive behaviour (ratio of attacks:displays) used among species(/experiment) and sexes. N = 29 female/ 46 male BB *M. zebra* (grey bars); 15 female/ 13 male *M. callainos* (blue and white bars). **a)** On average *M. callainos* were significantly more aggressive (p < 2e-16), and there is no difference in the total amount of aggressive behaviour from females and males (p = 0.939). **b)** *M. zebra* are only significantly less aggressive in the absence of a conspecific stimulus fish (Exp. 3, p < 2e-16). N = Exp. 1, 12 *M. zebra*; Exp. 2, 13 *M. callainos*; Exp. 3 12. *M. zebra*. **c)** The attack:display ratio does not differ significantly between species (p = 0.089), but males display more frequently than females which use a higher proportion of attacks (p = 9.45e-05).

#### Aggression biases in pairwise intruder choice tests

As expected, *M. zebra* males display significantly more aggressive behaviour towards conspecifics when given the choice of BB conspecific and B heterospecific stimulus males (glm; z_1,19_, = -16.03, p <2e-16, fig. 5, Exp 1).

**Figure 5.**
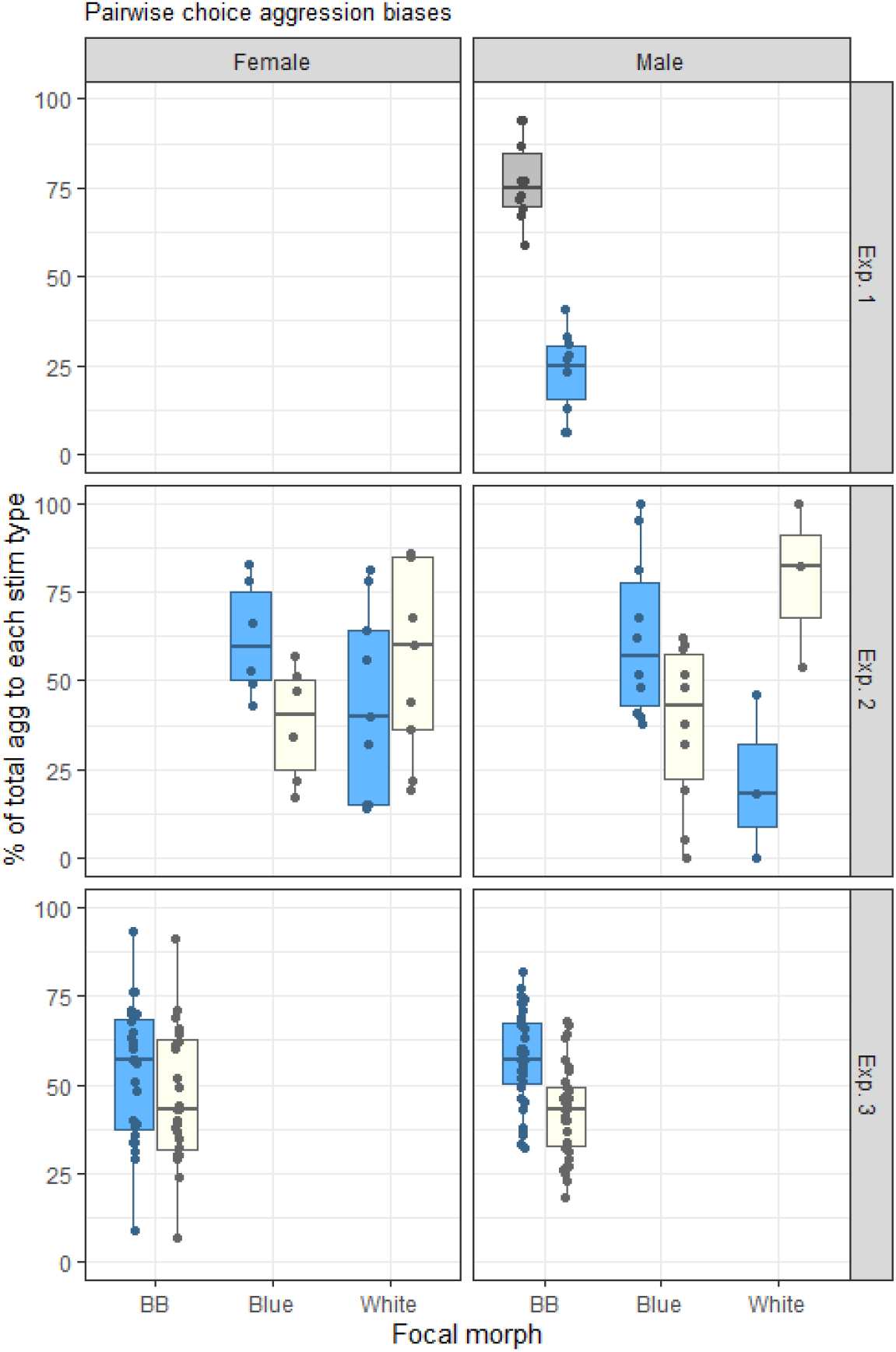
In laboratory based aggression trials: Exp.1) As expected, *M. zebra* males bias aggression towards conspecifics (n = 10, p < 2e-16). Exp. 2) Both *M. callainos* morphs showed a significant tendency to bias aggression towards intruders of the same morph as themselves (p < 2e-16). Exp. 3) Overall the *M. callainos* B morph stimulus fish received significantly more aggression from heterospecific *M. zebra* than the W morph (p < 2e-16).

Within *M. callainos*, the interaction between colour morph of focal and stimulus fish significantly affects proportion of aggression received, while sex has no effect (z_1,55_ Interaction = 12.48, p < 2e-16, Sex = 0.00, p = 1.00, fig. 5, Exp 2). To further clarify whether the difference in the proportion of aggression directed to stimulus fish of each colour morph is due to an own morph aggression bias (*i*.*e*. each colour morph is more aggressive to other males of the same colour) or an overall common morph aggression bias (*i*.*e*. males of both colour morph direct more aggression towards males of the common colour morph), stimulus type was recoded from Blue/White to either Other/Own or Common/Rare: Own-morph stimulus fish received significantly greater proportion of aggression than the other-morph stimulus fish, common morph stimulus fish did not receive a greater proportion of aggression overall in this experiment (z_1,55_ Other/Own z = 12.48, p < 2e-16, Common/Rare z = -0.30, p = 0.768).

When *M. zebra* were presented with the choice of *M. callainos* B and W stimulus pairs, overall both females and males preferentially attacked the common B morph (z_1,127_ Stim morph = -8.79, p < 2e-16, Sex = 0.01, p = 0.991, fig. 5, Exp 3). However, the level of aggression (total aggression count to both morphs) and the strength of aggression bias (proportion of aggression to B morph) differs between populations and sexes: *M. zebra* from Nkhata Bay (NB) and Thumbi West (TW) were significantly more aggressive to the heterospecific stimulus fish overall than those from Chiofu Bay (CB), and females overall were more aggressive than males (glm; z_2,63_ NB = 3.36, p = 0.0008, TW = 9.68, p < 2e-16, Sex = -2.62, p = 0.009, fig. 6a). In regards to strength of heterospecifc common morph aggression bias, we found that NB and TW focal fish displayed a significantly lower proportion of aggression towards the blue morph than those from CB (*i*.*e*. a weaker bias), and that although females were more aggressive overall, they also showed a weaker common morph aggression bias than males (glm; z_2,63_ NB = -2.50, p = 0.012, TW = -4.82, p = 1.47e-06, Sex = 2.05, p = 0.041, fig. 6b).

**Figure 6.**
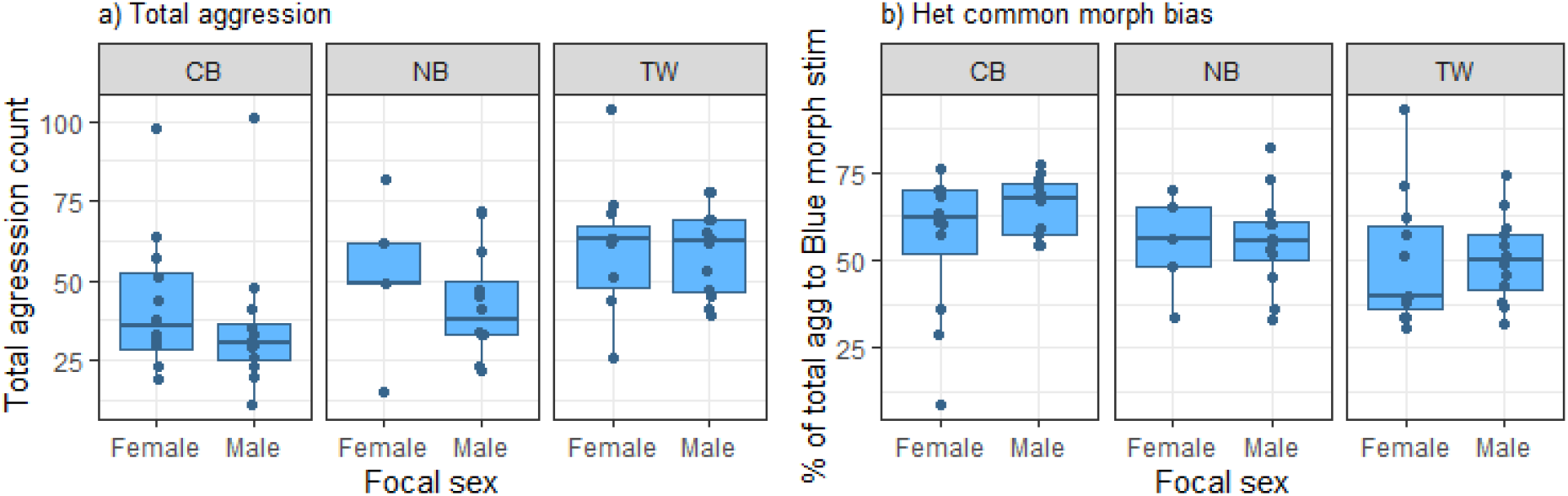
Differences in levels of aggression and aggression biases depending on *M. zebra* source population and sex (n females/males = CB 12/12, TW 5/12, NB 12/12). **a)** *M. zebra* from CB were significantly less aggressive in these experiments (compared to NB p = 0.0008, TW p < 2e-16), and females more aggressive overall than males (p = 0.009). **b)** *M. zebra* from CB showed a significantly stronger common (B) morph aggression bias than those from NB (p = 0.012) and TW (p = 1.47e-06), and females also displayed a weaker aggression bias than males (p = 0.041).

## Discussion

Our field observations of *Maylandia callainos*, a polychromatic mbuna cichlid from Lake Malawi, indicated that common (blue) morph territorial intruders received more aggression than rare (white) morph intruders.

Pairwise intruder choice tests in a controlled laboratory setting demonstrated that males and females of each morph bias aggression towards their own morph. These results add to the evidence that inherent own-morph aggression biases, which result in negative frequency dependent selection on rare colour morphs, can

contribute to balancing selection and thereby promote the maintenance of polychromatism (Dijkstra *et al*. 2008; Lehtonen 2014; Scali *et al*. 2020).

While this and previous studies (*e*.*g*. Dijkstra *et al*. 2008; Dijkstra *et al*. 2009a; Seehausen & Schluter 2004; Lehtonen 2014; Scali *et al*. 2020) have shown that aggression biases can be involved in stabilising polychromatism, to our knowledge this is the first study to identify benefits to rare morph cichlids which may result from receiving less aggression in the natural environment. We found there to be significantly more territory holding rare morph males than would be expected given the proportions of the two colours in the general population. Our observations suggest that the over-representation of rare morph territory owners may be influenced by two factors. Firstly, blue and white males appear to have higher tolerance of each other as neighbours, being found on average significantly closer to each other than blue morph individuals. Secondly, both territorial and non-territorial white morph individuals spend more time feeding, which suggests that the rare colour morph may benefit from lack of recognition during competition for non-mating related resources (Dijkstra *et al*. 2008; Lehtonen 2014; Pérez i de Lanuza G *et al*. 2017; Scali *et al*. 2020).

Further to showing that rare morph individuals can benefit from reduced intraspecific aggression, we found that a closely related ecologically similar heterospecific (*Maylandia zebra*) also biases aggression towards the *M. callainos* blue morph. While these results are in conflict with those from another cichlid fish system, which suggest that rare morphs may be disadvantaged by greater heterospecific aggression (Lehtonen *et al*. 2015), given that aggression among heterospecifics is often higher among more similar coloured than more differently coloured species (Genner *et al*. 1999; Pauers *et al*. 2008; Anderson & Grether 2010; Losin *et al*. 2016), it is not surprising to find that in some cases rare morph individuals may receives less aggression from a heterospecific which is more similar in colour to the common morph. We also found, however, that although heterospecific females were more aggressive overall, they also showed a weaker blue morph aggression bias than males. We speculate that a lower level of discrimination among morphs by heterospecific females, and the greater use of direct attacks compared to display behaviours (this study and Arnott & Elwood 2009), may be due to the difference in competition among females and males (*i*.*e*. greater heterospecific competition among females for access to shelters among the rocks during incubation of offspring).

In cichlids and other taxa, laboratory studies have shown that in species which differ in colour among allopatric populations, males tend to bias aggression towards males from their own population (Tyers & Turner 2013; Bolnick *et al*. 2016; Cooke & Turner 2018; Yang *et al*. 2018). In this study, we found that heterospecific aggression also varies depending on whether a pair of species occurs in sympatry or allopatry. The level of aggression (total aggression count to both *M. callainos* morphs) differs between *M. zebra* populations: *M. zebra* from Nkhata Bay (NB) and Thumbi West (TW), which co-occur with *M. callainos*, were significantly more aggressive to *M. callainos* than those from Chiofu Bay (CB), which are naïve to *M. callainos*. These findings support the hypothesis that aggression among heterospecifics may often not simply be due to misdirected aggression among species (Peiman & Robinson 2010), which would be indicated by higher levels of aggression from the allopatric *M. zebra* population (CB). The persistence of heterospecific aggression at NB support the idea that it has an adaptive function in long-term co-existing multi-species communities (Peiman & Robinson 2010; Losin *et al*. 2016). Although there are no *M. callainos* at Chiofu Bay, this location is home to another closely-related species (*M. esterae*) which has blue males, and brown, orange and orange blotch females. *M. zebra* at Chiofu Bay therefore do co-occur with a similar blue morph fish, but no white morph fish and we found that the *M. zebra* from this location has a stronger blue-morph aggression bias than the other *M. zebra* populations which co-occur with blue and white *M. callainos*. A previous study of a polymorphic frog species found stronger aggression biases among morphs when they occur in allopatry compared to when they are found in sympatry (Yang *et al*. 2018). Our results show a similar pattern in heterospecific aggression; a weaker blue morph aggression bias in *M. zebra* populations which coexist with both colour morphs.

Our results indicate that a rare colour morph may benefit from lack of recognition as a resource competitor, by both conspecifics and heterospecifics. This results in rare morph individuals receiving less aggression and gaining improved access to territories and food. This can benefit rare morph individuals while they are rare, but then what prevents them from increasing in frequency until fixation? Firstly, we found that rare (white) morph individuals were more aggressive towards their own morph than they were to the common (blue) morph, which would result in white morph individuals experiencing increasing levels of aggression as they became more common. Secondly, the lower level of heterospecific aggression bias towards the common morph, in populations with blue and white morphs, suggests that heterospecifics learn or evolve the ability to recognise rare morph individuals as competitors. The ability to recognise rare morph individuals may increase as they become more common: TW has the highest frequency of white morph individuals and the weakest common morph aggression bias by heterospecifics. Finally, female preference for common-morph males may result in a

disadvantage to rare morph males (Roberts *et al*. 2009). The genes responsible for the expression of the melanin-disrupted (“blotched”) morphs are almost always closely linked to a dominant female determiner, and so are generally much more common in females. This suggests that these colour phenotypes are disadvantageous to males, although they may be advantageous to females by providing increased crypsis or reduced aggression from conspecifics and/or heterospecifics.

## Conclusions

Our results support previous studies indicating that negative frequency-dependent antagonism can be generated by own-morph aggression biases among conspecifics in cichlids which display polychromatism. We find that heterospecifics show reduced aggression to rare morph individuals, suggesting that heterospecific aggression may also facilitate invasion of rare colour morphs into a population. We identify potential advantages to rare morph individuals in the field, in terms of territory and foraging.

## Acknowledgements

This work was supported by Bangor University PhD scholarships to AMT and GMC. We thank Aaron Scott, Winnie Courtene-Jones, Beth Powell for help with field recording, and James Whitman for collecting live fish. We are grateful to the Department for National Parks and Wildlife in Malawi for allowing us to collect fish from Lake Malawi National Park. All behavioural experiments were approved by Bangor University ethical review. Thanks also to P.D.Dijkstra for and another anonymous reviewer for comments on an earlier version of this manuscript.

## Notes

### Competing Interest Statement

The authors have declared no competing interest.

